# Hierarchical organization of rhesus macaque behavior

**DOI:** 10.1101/2021.11.15.468721

**Authors:** Benjamin Voloh, Benjamin R. Eisenreich, David J-N. Maisson, R. Becket Ebitz, Hyun Soo Park, Benjamin Y. Hayden, Jan Zimmermann

## Abstract

Primatologists, psychologists and neuroscientists have long hypothesized that primate behavior is highly structured. However, fully delineating that structure has been impossible due to the difficulties of precision behavioral tracking. Here we analyzed a dataset consisting of continuous measures of the 3D position of fifteen body landmarks from two male rhesus macaques (*Macaca mulatta*) performing three different tasks in a large unrestrained environment over many hours. Using an unsupervised embedding approach on the tracked joints, we identified commonly repeated pose patterns, which we call postures. We found that macaques’ behavior is characterized by 49 distinct identifiable postures, lasting an average of 0.6 seconds each. We found evidence that behavior is hierarchically organized, in that transitions between poses tend to occur within larger modules, which correspond to intuitively identifiably actions; these actions are in turn organized hierarchically. Our behavioral decomposition allows us to identify universal (cross-individual and cross-task) and unique (specific to each individual and task) principles of behavior. These results demonstrate the hierarchical nature of primate behavior and provide a method for the automated “ethogramming” of primate behavior.

## INTRODUCTION

Understanding the principles behind the organization of behavior has long been an important problem to ethology, psychology, and neuroscience (Krakauer et al., 2017; Tinbergen, 1951; Gallistel, 2013; Anderson and Perona, 2014; Calhoun and El Hady, 2021; Periera et al., 2020). Macaques are especially important in this regard because of their pivotal role as a model organism for biomedical research (Rudebeck et al., 2019; Buffalo et al., 2019). Indeed, a great deal of research has benefited from the rudimentary tracking and identification of behavior in laboratory tasks in macaques. However, precise measurement of behavior has generally been limited to a single motor modality (typically the eyes or arm) under conditions of bodily constraint. As a result, we have an impoverished understanding of behavior in the natural context, involving the free movement of full bodies in three-dimensional space (Hayden et al., 2021).

Recent years have seen a great deal of success in the development of camera-based systems for tracking the behavior of small animals, including worms, flies, and mice (Mathis and Mathis, 2020; Periera et al., 2020; Calhoun et al., 2019; Sturman et al., 2020; Hsu and Yttri, 2020; Bohnslav et al., 2021). There is now growing interest in larger species, including primates (Marks et al, 2021; Dunn et al., 2021; Bain et al., 2021). These tracking systems have allowed for the automated identification of specific meaningful behavioral units (“ethogramming”) in these species (Marshall et al., 2021; Berman et al., 2016; Wiltschko et al., 2015; Bain et al., 2021). Results of these analyses have shown that behavior in these organisms consists of simple motifs that are repeated and are organized into a hierarchical structure (Berman et al., 2016; Marshall et al., 2020). These methods are important because they can provide quantitative answers to longstanding questions at the core of behavioral science. However, we do not know whether these principles hold true for larger animals with more complicated behavioral repertoires. In particular, macaque behaviors might not obey the same principles because their bodies have much higher degrees of freedom, and consequently their behavior has much higher dimensionality (Bala et al., 2020).

Our laboratory has developed a system that can perform detailed three-dimensional behavioral tracking in rhesus macaques with high spatial and temporal precision (Bala et al., 2020 and 2021). Our system uses 62 cameras positioned around a specially designed open field environment (2.45 × 2.45 × 2.75 m) in which macaque subjects can move freely in three dimensions and interact with computerized feeders (haydenlab.com/tracking). We used this system to track the position of 15 joints at high temporal and spatial resolution as our subjects performed three different behavioral tasks. The data collected by this system open up the possibility of automated behavioral identification and analysis in macaques.

Previous unsupervised approaches to quantifying behavior were centered on actions (Marshall et al., 2021; Berman et al., 2016). In contrast, our approach starts with the configuration of landmarks (“postures”) as the fundamental unit of behavior. Specifically, we generated 23 variables corresponding to the angles between all major joint pairs and the velocity of the subject in three dimensions. We then performed dimensionality reduction to identify postures, and graph theoretic methods to identify extended actions. We find that behavior naturally clusters into 49 distinct postures. Further graph-theoretic analyses show that postures are organized into specific actions. These actions correspond to nameable, intuitive behaviors, and are further organized into higher categories. Together, these results confirm that monkey behavior obeys the same hierarchical organizational principles that simpler organisms do. These results also indicate that our pipeline can overcome the daunting problems faced by the high dimensionality of movement in monkeys.

We examined behavior of two macaque subjects performing one of two different tasks, or, in a third condition, no task. This design allowed us to examine the effect of task and of individual on the organization of behavior. We found prominent cross-individual differences and only modest cross-task differences. Moreover, we found that the composition of behavior (as inferred by adjusted mutual information) is more stable during task performance than during task-free behavior. This finding demonstrates changes in the way behavioral repertoires are selected on the basis of behavioral context. Overall, these findings demonstrate that it is possible to obtain automatic behavioral ethograms in macaque monkeys and delineate the organization of behavior across contexts in this important species.

## RESULTS

We studied the behavior of two rhesus macaques under three different experimental conditions (see **Methods)** in a large open cage that allowed for free unimpeded movement (a 2.45 × 2.45 × 2.75 m cage with barrels, **Figure 1A**, Bala et al., 2020). Each subject performed under one of three possible task conditions per day (see below and **Methods**). Each daily session took about 2 hours. Each task condition was repeated three times over three different days (fully randomized and interleaved order). Our dataset therefore consists of 18 sessions (9 for each subject; divided into two different task conditions, 6 task-OFF, 12 task-ON) for a total of 31.4 hours, or about 3.3 million frames. Behavior was tracked with 62 high-resolution machine vision video cameras, and pose (3D position of 15 cardinal landmarks, see **Methods**) of each macaque was determined using OpenMonkeyStudio (Bala et al., 2020, **Figure 1B**) with secondary landmark augmentation (Bala et al., 2021).

**Figure 1.**
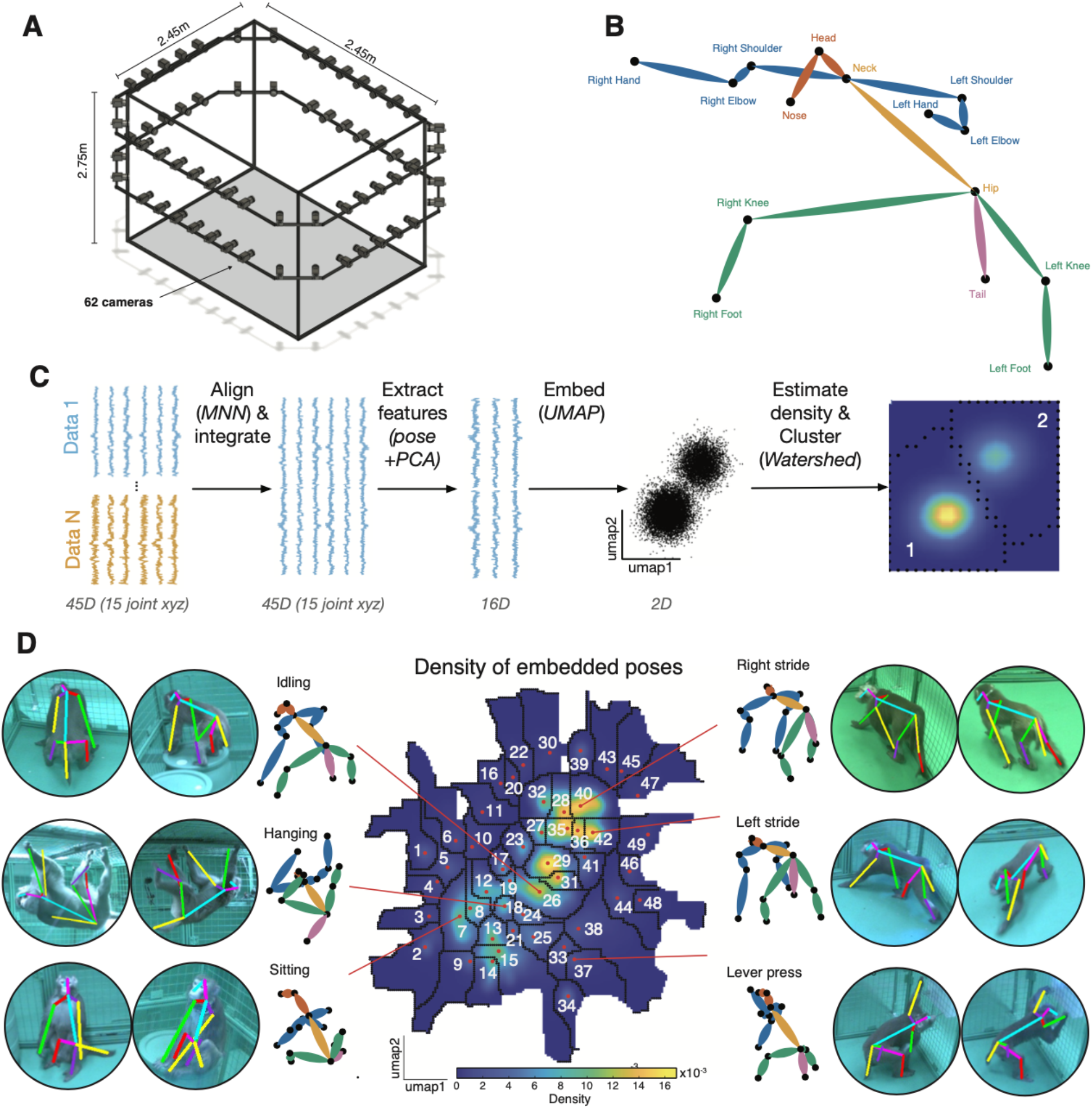
Identification of postures in an open-field environment. **(A)** Depiction of the cage environment. 62 cameras were mounted on an exoskeleton, facing inwards. **(B)** Reconstructed pose was defined by 15 landmarks. **(C)** Outline of general methodological approach. See **Methods** for details. **(D)** The heatmap denotes the density of embedded samples. Select postures are visualized here, both as the mean posture within clusters (monkey stick figures) and example reprojections onto the raw data.

### Embedding of macaque posture results in semantically meaningful clusters

We developed a novel pipeline to characterize behavioral states based on tracked poses. Our pipeline is a variation of one developed by Berman and colleagues to characterize the behavior of flies (**Methods**, **Figure 1C**, Berman et al., 2014 and 2016). The major difference is with the way that pose data were structured at the beginning of the pipeline. Briefly, poses were translated using the neck as the reference. Then, the pose of the subject in each individual frame was rotated to face a common direction (see **Methods**). This rotation was defined via two vectors corresponding to the spine and shoulders. Next, poses were size-scaled (with size defined as hip to neck distance) so that subjects matched. This process produced normalized postural orientations. Finally, to further reduce individual variation in poses, we aligned poses of individual subjects via a variation of the Mutual Nearest Neighbors approach (a local alignment procedure; see **Methods** and Haghverdi et al., 2018).

After normalization, we embedded poses using all collected data to generate a single overall postural embedding. We used a dimensionality reduction technique known as uniform manifold approximation and projection (UMAP, McInnes et al., 2018). This process results in a reprojection onto two dimensions in which similar poses are adjacent in the resulting low-dimensional space. We then performed a kernel density estimation to approximate the probability density of embedded poses at equally interspersed points. The color in the resulting plot reflects the probability of each pose in our dataset (**Figure 1C and D**).

Inspection of this postural embedding reveals a clear clustered organization. Each cluster reflects a set of similar poses that are relatively distinct from other sets of poses. To formally identify these clusters, we used the *watershed algorithm* on the inverse of the density map (Berman et al 2014). This algorithm treats each peak as a sink and draws boundaries along lines that separate distinct basins. We found that the resulting *embedding space* contains 49 distinct clusters. These clusters correspond to sets of closely related poses (**Figure 1D**). We verified that the embedding space captures differences in poses by correlating the euclidean distance of pose features with that of embedded points (bootstrap test, mean Pearson *r* = 0.45, p < 0.001). We refer to the clusters of poses as *postures*. Visual inspection reveals that these postural states are semantically meaningful in the sense that they correspond to recognizable postures such as left or right stride, sitting, hanging etc. Each posture lasted on average 0.612 ± 0.0015 (s.e.m) sec. The clustered nature of this embedding space confirms that macaque behavior is composed of stereotypical postures.

### Directed graph analysis of posture transitions reveals behavioral modularity

We next sought to understand how postures combine to form recognizable behaviors. To do this, we sought to identify sets of postures with a high probability of occurring in sequence. We therefore computed *transition probability matrices* for the specific postures identified above. Any organization in sequences of postures will show up in the form of increased likelihood of specific transitions between the postures within the sequence. Our goal, then, is to discover sets of postures that have a high probability of transitioning between one another, but not other sets of postures.

The transition probability matrix, in graph theoretic terms, is a directed graph. In this framework, nodes that form strong links between each other are referred to as *modules* or *communities*. Because of this, we refer to sets of postures that have a high probability of co-transition as “behavioral modules”. These modules are roughly equivalent to what are sometimes called actions (Anderson and Perona, 2014). The identification of these modules allows us to re-sort the transition probability matrix such that there are blocks on the diagonal; these blocks correspond to the behavioral modules.

To formally identify behavioral modules in the transition matrix, we used a recently-developed algorithm named *Paris* (Bonald et al., 2018). This algorithm performs hierarchical clustering on the graph derived from the transition matrix and returns a tree describing the distance between poses and their composing modules (we will return to examine the hierarchical structure of behavior below). Next, to determine the optimal number of behavioral modules, we proceed to cut the tree at a series of hierarchical levels and compute a modularity score for each cut tree. We then choose the cut that corresponds to the tree with the maximal modularity score. The modules that result from this cut give the highest average within-module posture-transition probability and the lowest average across-module posture-transition probability. Formally speaking, they maximize the difference between these two measures. From this process we can identify the most likely behavioral modules.

A transition matrix from an example *task-OFF* session is depicted in **Figure 2Ai**. For this session, the number of modules that maximizes the modularity score is 5, indicating that the best fitting classification has 5 discrete behavioural modules (**Figure 2Aii**). As illustrated in the figure, these behavioral modules hew closely to nameable actions (**FIgure 2B)** such as walking (**Video 1**), swaying (**Video 2**), climbing (**Video 3**), jumping (**Video 4**), and idling (**Video 5**). The modular nature of the behaviors in this session is clearly visible in the sorted transition matrix (**Figure 2Aiii**).

**Figure 2.**
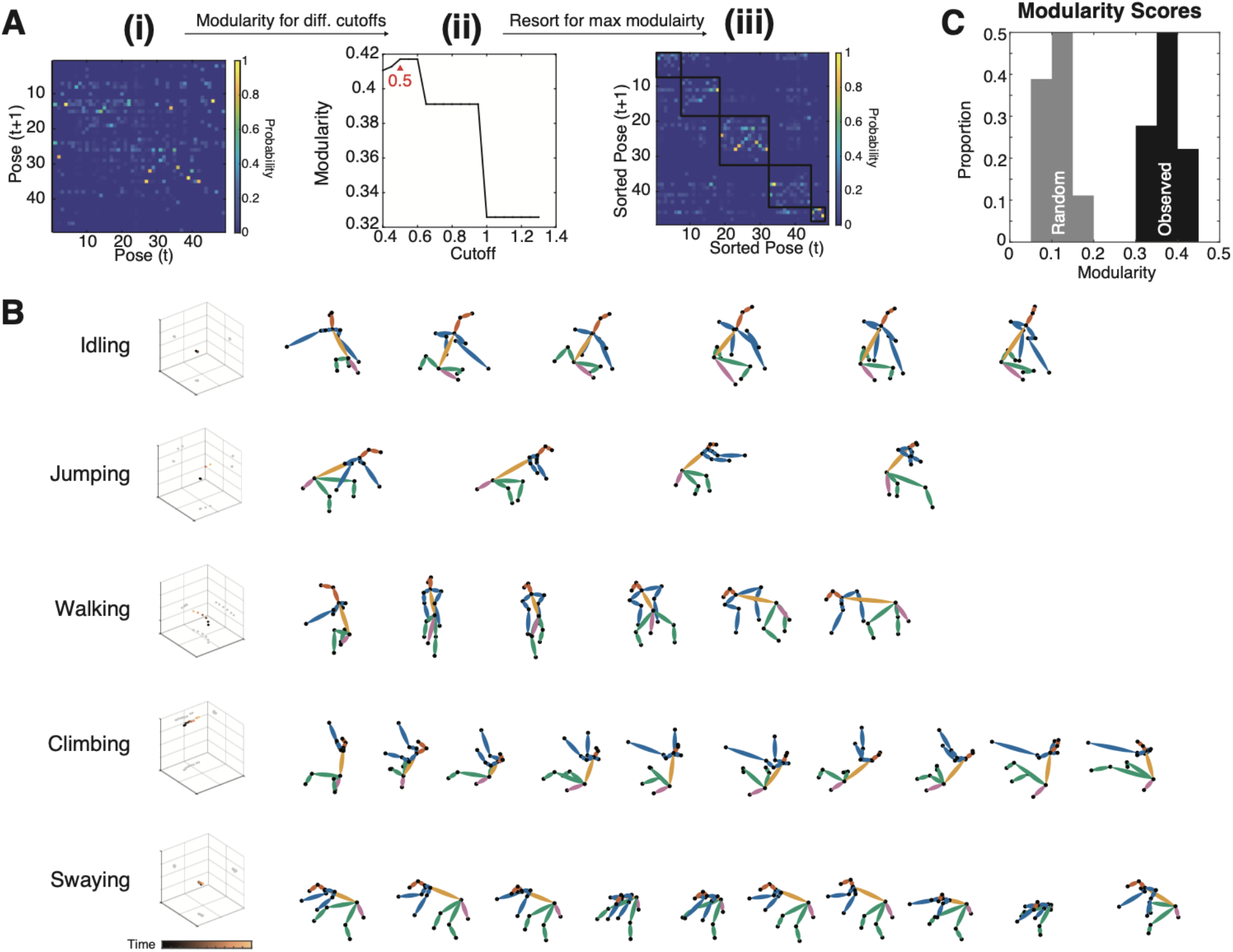
Postures are organized into behavioral modules. **(A)** Example of modularity in one dataset. **(i)** The original transition probability matrix. **(ii)** The modularity score for multiple different cutoffs of the dendrogram **(iii)** Same as (i) but sorted according to the results of (ii). Now, it is evident that transitions between poses occur within modules (highlighted with black squares). **(B)** All modules in this example. These correspond to semantically meaningful sequences of poses. **(C)** Histogram of the maximal modularity score, both for observed (dark) and randomized (light) transition matrices. Modularity is higher than expected by chance.

We tested for modularity by computing the Modularity Score. All 18 of the individual datasets we collected individually showed statistically significant evidence of modularity (randomization test, p<0.001, **Figure 3C**). The average number of unique behavioral modules in each session was 3.8 ± 0.15 (sem), and each behavioral module lasted 2.73 ± 0.0327 sec. Across all sessions, the duration of modules ranged from 0.47 - 16.8 sec. Together, these results indicate that subjects’ behavior is organized into discrete behavioral actions that consist of stereotyped patterns of postures.

**Figure 3.**
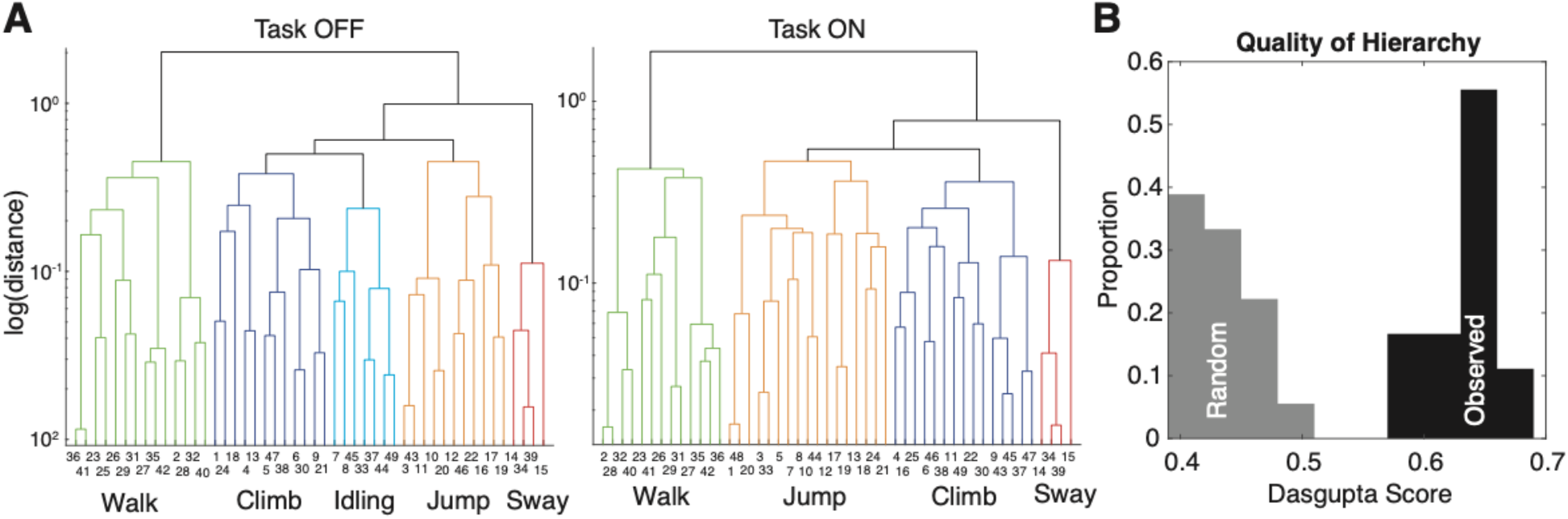
Pose transitions are hierarchically organized. **(A)** Dendrograms for two example sessions from the same individual, for a task OFF (left) and task ON (right) condition. **(B)** Dasgupta score, measuring hierarchical organization, from transition matrices derived from observed (dark) and randomized (light) transitions.

### Transitions are hierarchically organized

Our next analysis investigated the hierarchical organization of behavior (**Figure 3)**. Dendrograms in this figure show the hierarchical organization of postures according to their transition probabilities. Higher level connections in this dendrogram show how different sets of poses are related. Not only are different behavioral modules recognizable actions (see above), but their relationship in the tree reveals this subject’s idiosyncrasies; for example, idling before climbing (**Figure 3A**). Moreover, across sessions, similar behavioral modules were composed of similar postures, highlighting the stable behavioral repertoire of one subject (more on this below). To quantify the degree of hierarchical organization, we calculated the Dasgupta score on these dendrograms, which quantifies the quality of hierarchical clustering on a graph (Dasgupta, 2016). A Dasgupta score above chance indicates that the observed tree indeed has connected components that are related to one another (Dasgupta, 2016). The Dasgupta score was significantly above chance for all 18 datasets (**Figure 3B;** randomization test, p<0.001).

### Behavioral organization is evident for lagged transitions

The previous sections indicate that at short timescales, behavior is modular and hierarchical. We next investigated the possibility of levels of organization defined by even longer timescales. To this end, we performed the same modularity analysis as above, but constructed the transition probability matrix with lags of up to 1000 transitions. Thus, if there are long timescale drivers of behavior, this should again be evident as above-chance behavioral modularity.

For the same *task OFF* example dataset as above, the modularity score decreases as a function of transition lag, before plateauing close to (but greater than) zero around a lag of about 100 transitions (**Figure 4Ai**). This result demonstrates that behavior is non-stochastic even on very long timescales, but that its organization decreases with timescale in a systematic and lawful way. This modularity is also reflected in the transition matrices for shorter, rather than longer, lags (**Figure 4Aii**). Across all 18 datasets, the average modularity shows steady decay up to ~100 transitions into the future (~60 sec), before plateauing close to chance levels (**Figure 4B**). This pattern suggests that not only do poses tend to co-occur in distinct behavioral modules, but that this organization is evident even when considering longer timescales.

**Figure 4.**
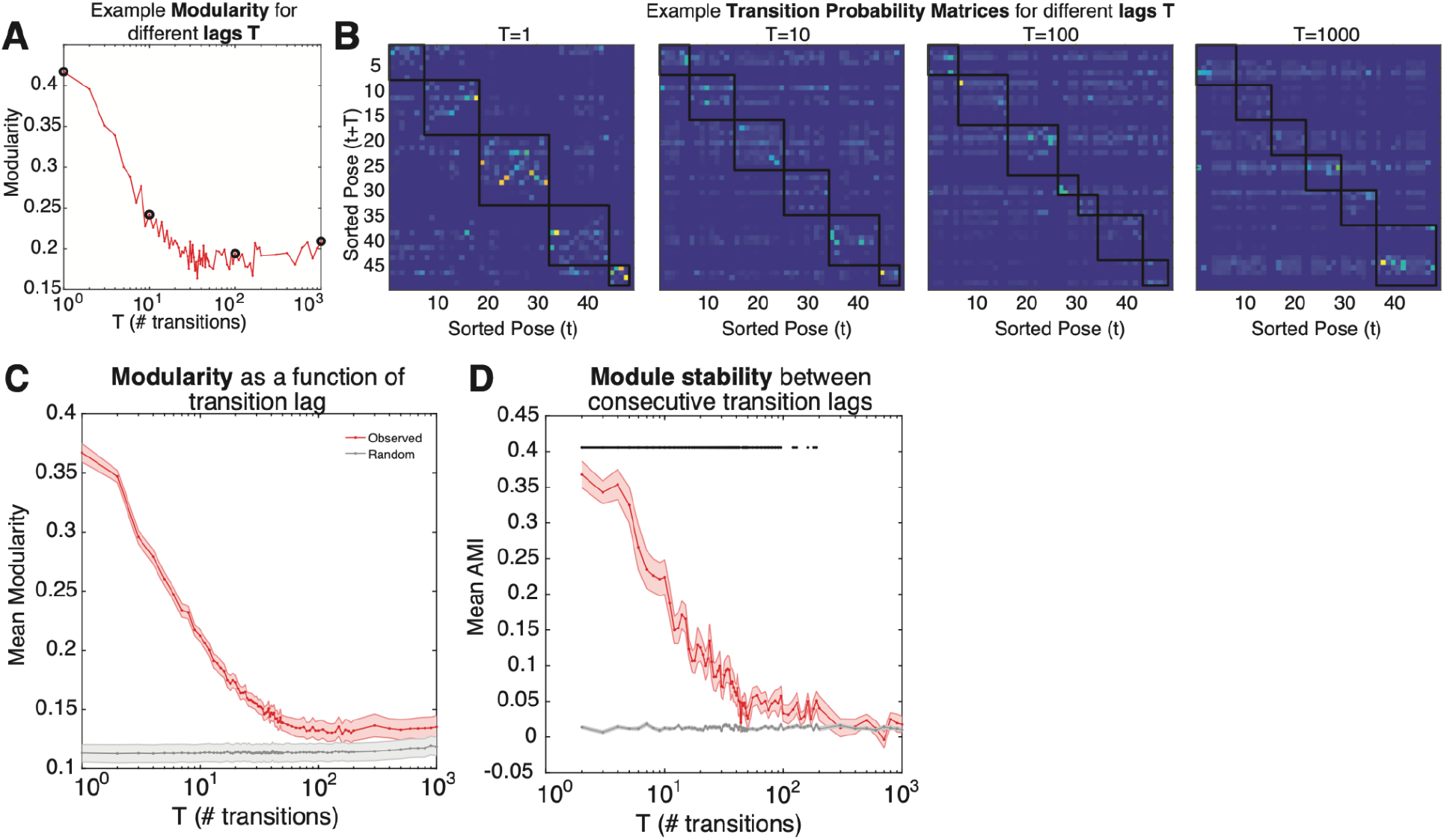
Modular and hierarchical organization of behaviour is evident for long timescales. **(A)** Example transition matrices and associated modularity (i) Transition matrices with lag 1, 10, 100, 1000. Note that states have been reordered according to module classification. (ii) Modularity as a function of transition lag. Black circles correspond to transition matrices in (i). **(B)** Mean modularity across all datasets, for observed (red) and randomized (grey) transition matrices. **(C)** Mean module stability (adjusted mutual information score) of module assignments between consecutive transition lags, for observed (red) and randomized (grey) transition matrices.

Because modules are computed independently for each transition matrix of different lags, a critical question is whether the extracted behavioral modules are consistent across transition lags. We assessed this by comparing module assignments using the *adjusted Mutual Information Score* (AMI; Vinh et al 2010; *see* **Methods**) between modules derived from transition matrices of consecutive lags. We found that cluster assignment was stable across transition lags (**Figure 4C**), up to ~100 transitions into the future (p<0.05, multiple comparison corrected). This finding is reassuring; it suggests that behavioral modules are composed of similar poses for as long as 100 transitions into the future. Taken together, these results indicate that current states hold information about states up to at least 100 transitions into the future.

### Variability in behavioral repertoire is driven by individual and task differences

Up to this point, we have considered the organization of behavior across all individuals and datasets, leaving open the question of how individual and task variability affects behavioral organization. To this end, we first determined if behavioral modularity - derived from transition probabilities with a lag of 1 - was affected by task and individual. We found that the modularity score varied as a function of individual (2-way ANOVA; F=8.08, p=0.013), but not task (F=0.14, p=0.87). Similarly, the Dasgupta score varied by individual (2-way ANOVA; F=5.2, p=0.0.39) but not task (F=0.1, p=0.90). Taken together, these results suggest that the degree of behavioral organization at the shortest timescale is driven by individuals but not environmental constraints.

While the degree of organization may not vary by task or individual, it is still possible that the composition of behavior differs. To this end, we compared module stability between individuals and tasks (**Figure 5)**. We computed the AMI between all pairs of datasets and asked if stability between pairs differed as a function of subject or task. We found that both individual and task influenced the degree of stability between pairs of datasets (**Figure 5A;** 2-way ANOVA; F(individual)=43.7, p<0.0001; F(task)=7.14, p=0.001). All individual/task combinations were also more stable than chance level (**Figure 5A**; randomization test, p<0.001). We also found that within-subject stability was higher than between subject stability (**Figure 5B**; unpaired T-test, T=9.4, p<0.001), indicating that while there is a significant amount of overlap in behavioral modules, subjects still tend to perform specific actions in idiosyncratic ways.

**Figure 5.**
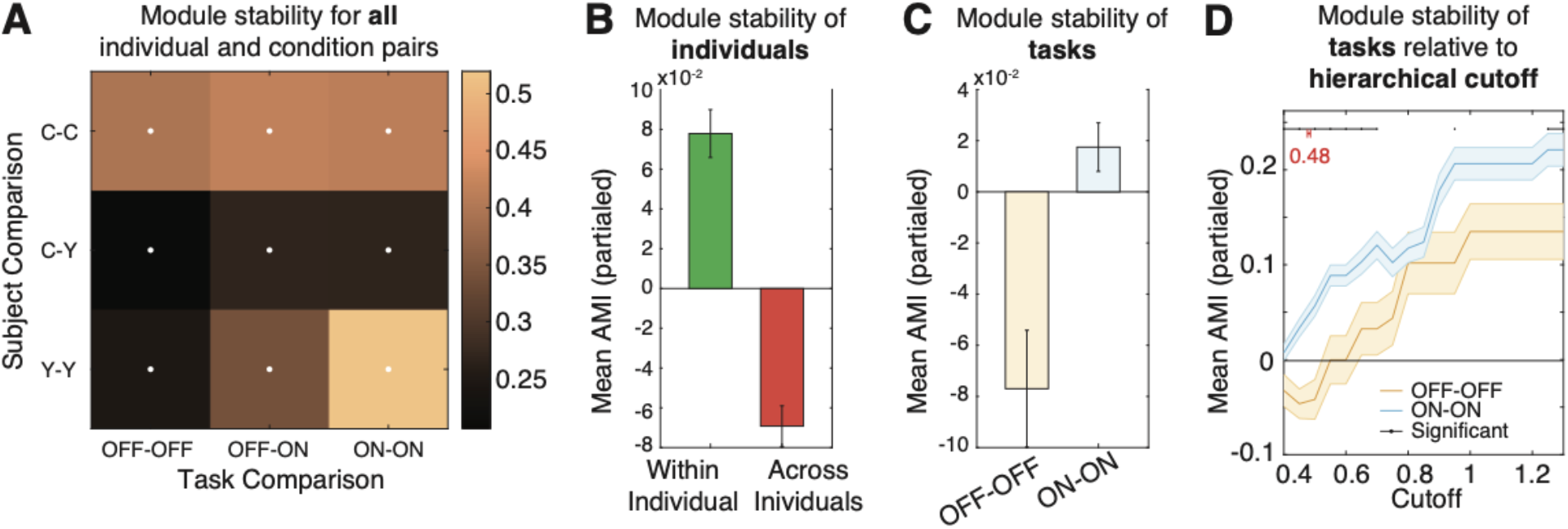
Behavioral modularity is universal and unique. **(A)** Mean module similarity (quantified via the adjusted Mutual Information (AMI) score) across datasets with the same or different subjects, and same or different task demands. White dots denote significant cells (randomization test, p<0.05, multiple comparison corrected). There is significant module overlap both between datasets of the same subject, and that of different subjects. **(B)** Comparison of module stability either within the same individuals (green) or across different individuals (between), where the effect of task has been partialled out. **(C)** Same as (B) but comparing module stability across different tasks, either task OFF-OFF (orange) or task ON-ON (blue) comparisons. The effect of individuals has been partialled out. **(D)** Mean+ SEM of module stability comparing task ON-ON (blue) or task OFF-OFF (orange) pairs, after partialling out the effect of individual. Black lines and dots denote significant differences (multiple comparisons corrected). The mean cutoff that maximized modularity is depicted in red.

We next asked if module stability varied as a function of task (**Figure 5C**). We found that sessions with a task were more similar to one another, rather than sessions with no task (unpaired T-test, T=4.16, p<0.001). This suggests that task demands are a strong constraint on the expression of behavior.

We further explored the effect of task on behavioral expression by quantifying stability for different cuts of the dendrograms associated with each dataset (**Figure 5D**). We found that *task ON* pairs showed more stable behavioral expression than *task OFF* pairs, and this was generally significant for lower and higher dendrogram cuts (unpaired T-test, multiple comparison corrected, p<0.05). Thus, environmental context constrains behavioral expressivity whether the span of individual behaviors is large or small.

### Timescale of behavioral organization is driven by individual, but not task, variation

We next asked if the timescales of behavioral organization differed by task and individual. We operationalized the notion of how many transitions into the future exhibited modular organization as the half-life of the function that relates modularity to transition lag (the modularity curve). Specifically, we fit an exponential model to the individual modularity curves, and determined the half-life associated with the exponent term (see **Methods)**. We found that modularity curves are well fit by the model (**Figure 6A inset**; mean adjusted R2=0.95 + 0.005). Half-lives varied significantly as a function of individual but not task (**Figure 6A**; 2-way ANOVA, F(individual)=26.8, p<0.001; F(task)=2.87, p=0.11). In other words, individuals varied in the extent of the temporal horizon of modular behavioral organization.

**Figure 6.**
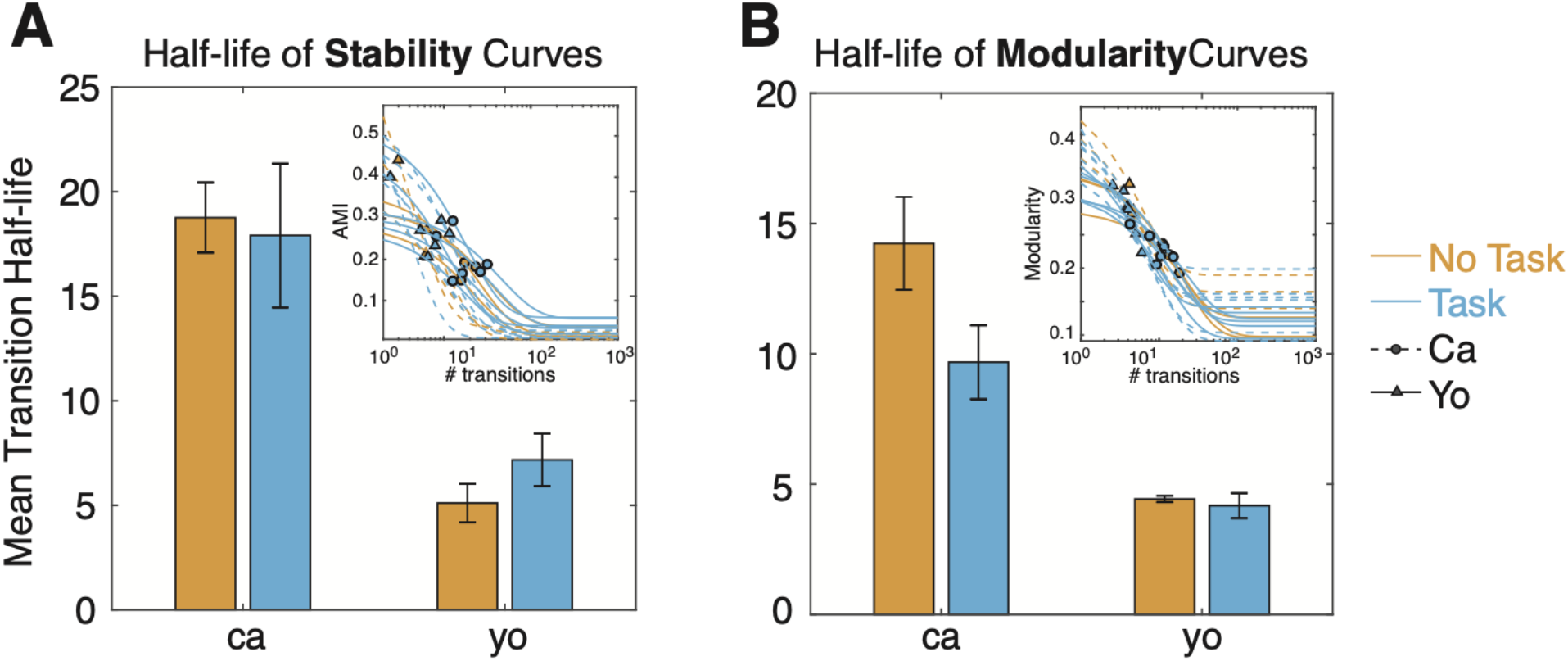
Timescale of behavioral organization varies as a function of individual, but not task. **(A)** Mean and standard error of half-life associated with fitted modularity curves. Fitted modularity curves are visualized in the inset, and plotted as a function of transition lags. Orange (blue) is for task OFF (ON) sessions, and solid (dotted) lines are for subject C (Y). The half-life of each curve is depicted as a solid circle (triangle), for subject C (Y). Mean half-life varies as a function of individual, but not task. **(B)** Same as (A) but calculated for AMI (i.e., module stability) curves. Mean half-life varies as a function of individual, but not task.

As noted above, modularity at different lags does not guarantee similar module composition. Thus, we repeated the same analysis as above, but fitting module stability curves (AMI as a function of transition lag) and extracted their half-lives (instead of from the modularity curves as before). Stability curves were well-fitted by the exponential model (**Figure 6B inset**; Mean adj. R2=0.68 + 0.02). Half-lives varied by individual, but not task (**Figure 6B**; ANOVA, F(individual)=18.3, p<0.001; F(task)=0.044, p=0.84)

## Discussion

Here we provide the first analyses of macaque behavior derived from quantitative 3D pose data. Our ability to perform these analyses relies on our recently developed OpenMonkeyStudio system, which allows for the tracking of major body landmarks as macaques move freely in a large space in three dimensions (Bala et al., 2020 and 2021). We find that, within the context of three different task conditions (including a no task condition), macaque behavior can be classified into 49 different *postures*, such as left and right strides, sitting, and hanging. We find that these postures in turn can be clustered into behavioral modules, such as walking, climbing, and swaying; these can in turn be organized into even higher-level structures. Thus, our method provides a hierarchical description of behavior that spans low-level postures and higher-level extended action sequences.

We find that these hierarchies vary both across individuals and task. Behavioral modules were more stable within the same individual than across the two individuals. In addition, the presence of a task resulted in more stable behavior composition as compared to when no task was present. Finally, we found that the timescale over which behavior was detectably organized varied strongly as a function of individual, but not task. Taken together, these results highlight the importance of both task demand and individual identity in determining the makeup of the hierarchy of actions, while also demonstrating that actions can have consistent cross-individual and cross-task properties. Our results also raise important lines of inquiry, including what factors may alter the timescale of structures behavior other than individual identity, identifying the extent to which the timescale of structured behavior depends on internally defined or externally imposed timescales, and identifying variation in individual actions that are relevant for task goals.

Multiple approaches exist to identify relevant behaviors based on pose data. These can, like our methods, rely on embedding pose features to discover low-level behaviors (Berman et al., 2016, Marshall et al., 2020), on fitting pose time series using Hidden Markov Models (Wiltschko et al., 2015; Calhoun et al., 2019), or on pre-trained, supervised neural network architectures (Marks et al., 2020; Sturman et al., 2019). Regardless of the method used, there are three important principles to consider that determine what inferences can be made to determine the structure of behavior. First, the timescale over which features are calculated determines the nature of the lowest level of behavior identified. In our case, because our model inputs correspond to instantaneous joint positions and speeds, our elementary unit of behavior is posture. Second, it is important to consider the way in which low-level behaviors are combined to form high-level actions. In our case, we do this solely on the basis of subsequent transitions, which allows for the discovery of actions with no strong a priori guess about their duration. Third, it can be constrained by previously identified behaviors, meaning that learning is semi-supervised (e.g., Marshall et al., 2021) or fully supervised (e.g., Marks et al., 2020; Struman et al 2019; Bain et al., 2021); we used an unsupervised method to identify behaviors. The unsupervised nature means our system can identify a much wider range of possible behaviors, including new ones not anticipated by existing theories.

The repertoire of behavior was more stable with an externally imposed task, suggesting that environmental demands may provide a force for behavioral stabilization. Stabilization can occur in one of two (not mutually exclusive) ways; either the repertoire of actions significantly shrinks during task, or actions themselves become less variable during task performance. Our data suggest the latter, as modules were stable even when we considered relatively high cutoffs of the dendrogram (which is to say, for larger and more-encompassing behavioral modules, Figure 5D). This is reminiscent of hunting behavior observed in zebra-fish placed in a prey-rich or prey-poor environment (Marques et al., 2020). In that study, animals exhibited behavioral motifs associated with exploitation and exploration, regardless of the environment. An intriguing future possibility direction would be to dissociate the sources of variation underlying variation in actions themselves, or which actions are expressed, on the basis of task demands.

One of the greatest potential benefits for statistical analysis of highly quantified behavior is in the prospect of automated ethogramming (Periera et al., 2020; Anderson and Perona, 2014; Hayden et al., 2021). By *ethogramming*, we mean the classification of pose sequences into specific behavior into ethologically meaningful categories such as walking, foraging, grooming, and sleeping (Hayden et al., 2021). Currently, constructing an ethogram requires the delineation of ethogrammatical category involves the time-consuming and careful annotation of behavior by highly trained human observers. Human-led ethogramming is slow, extremely costly, error-prone, and susceptible to characteristic biases (Anderson and Perona, 2014; Kardish et al., 2014; Holman et al., 2015; Tuyttens et al., 2014). For these reasons, it is simply impractical for even moderately large datasets, collected either in an open environment or the home cage (Womelsdorf et al., 2021; Hjunag et al., 2010). These kinds of datasets require automated alternatives. Automated ethogramming requires both high quality behavioral tracking and novel methods applied to tracked data that result in detection of meaningful categories. Such techniques have not, until recently, existed for primates. Our methods take the raw information needed for ethogramming - pose data - and infer posture and higher-level categories from it. As such, they provide the first step towards automated ethogramming in primates. We are particularly optimistic about the potential benefits of ethogramming for systems neuroscience. Relating behavior to neural circuits and networks is an important goal in the field, so being able to quantify behavior more rigorously – without sacrificing freedom of movement or naturalness – is likely to be invaluable for future studies.

## METHODS

### Animal Care

All research and animal care procedures were conducted in accordance with University of Minnesota Institutional Animal Care and Use Committee approval and in accord with National Institutes of Health standards for the care and use of non-human primates. Two male rhesus macaques served as subjects for the experiment. One of the subjects (C) had previously served as subjects on standard neuroeconomic tasks, including a set shifting task, a diet selection task, intertemporal choice tasks, and a gambling task (Ebitz et al., 2019; Farashahi et al., 2019; Blanchard et al., 2013 and 2014; Azab et al., 2018). This subject also participated in a study of foraging decision-making in the same environment as the current study (Eisenreich et al., 2019). The second subject (Y) was naïve to all laboratory tasks before training for this study. Both subjects were fed ad libitum and pair-housed with conspecifics within a light and temperature-controlled colony room.

### Behavioral Training and Tasks

Subjects were tested in a large cage (2.45 × 2.45 × 2.75 m) made from framed panels consisting of 5 cm wire mesh (Bala et al., 2020). Subjects were allowed to move freely within the cage in three dimensions. The wire mesh allowed them to climb the walls and ceiling, which they often did. Five 208 L drum barrels, weighted with sand, were placed within the cage to serve as perches for the subjects to climb and sit on. There was also a small, swinging tire hung from the centre of the ceiling of the cage. In sessions with a task, four juice feeders were placed on the wire mesh walls at each of the four corners of the cage. Feeders were placed at various heights, including atop barrels. The juice feeders consisted of a 16 × 16 LED screen, a lever, buzzer, a solenoid valve (Parker Instruments) and were controlled by an Arduino Uno microcontroller. Each feeder ran (the same) custom Arduino code.

We first introduced subjects to the large cage environment and allowed them to become comfortable in it. This process consisted of placing them within the large cage for progressively longer periods of time over the course of about five weeks. We monitored their behavior for signs of stress or anxiety. Notably, we did not observe these symptoms; indeed, subjects appeared to be eager to begin their sessions in the large cage, and somewhat reluctant to terminate them. Nonetheless, to ensure that the cage environment had positive associations, we provisioned the subjects with copious food rewards (chopped fruit and vegetables) placed throughout the environment. We then trained subjects to use the specially designed juice dispenser. We defined acquisition of task understanding as obtaining juice rewards in excess of their daily water minimum. For both subjects, acquisition of reliable lever pressing took about three weeks.

On any given day, animals performed one of three task conditions: (1) *a controlled depletion task*, (2) *a random depletion task*, and (3) *no task* (the same tasks were used in Eisenreich et al., 2019). In the no task condition, animals were free to explore the environment but no juice feeders were available. For both the controlled and random depletion tasks, each feeder was programmed to deliver a specific reward size on pressing of a lever; it started high and decreased by a specified amount. In the controlled condition each feeder delivered a base reward consisting of an initial 2 mL of juice that decreased by 0.125 mL with each subsequent delivery (turn). In the random condition, feeder depletion rates were the same as the controlled depletion condition. However, feeders randomly increased or decreased the juice delivery amount by 1 mL in addition to the base reward schedule at a probability of 50%. Both feeder types delivered rewards following their respective schedules until reaching the base value of 0, at which point the patch was depleted and no more rewards were delivered.

### Data acquisition

Images were captured with 62 cameras (Blackfly, FLIR), synchronized via a high-precision pulse generator (Aligent 33120A) at a rate of 30 Hz. The cameras were positioned to ensure coverage of the entire arena, and specifically, so that at least 10 cameras captured the subject with high-enough resolution for subsequent pose reconstruction, regardless of the subject’s position and pose. Images were streamed to one of 6 dedicated Linux machines. The entire system produced about six TB of data for a two hour session. After data acquisition, the data were copied to an external drive for processing on a dedicated Linux server (Lambda Labs).

To calibrate the camera’s geometries for pose reconstruction, a standard recording session began with a camera calibration procedure. A facade of complex and non-repeating visual patterns (mixed art works and comic strips) was wrapped around two columns of barrels placed at the centre of the room, and images of this calibration scene were taken from all 62 cameras. These images were used to calibrate the camera geometry (see below). This setup was then taken down, and the experiment began.

### Pose reconstruction

We first extracted parameters relating to the cameras’ geometry for the session. To this end, we used a standard structure-from-motion algorithm (*colmap;* Schonberg and Frahm, 2016) to reconstruct the space containing the 3D calibration object and 62 cameras from the calibration images, as well as determine intrinsic and extrinsic camera parameters. We first prepared images by subtracting the background from each image in order to isolate the subject’s body. Then, 3D center-of-mass trajectories were determined via random sample consensus (RANSAC). Finally, the 3D movement and subtracted images were used to select and generate a set of maximally informative cropped images, such that the subject’s entire body was encompassed. To reduce the chance that the tire swing would bias pose estimation, we defined a mask of pixels to ignore that encompassed the tire’s swinging radius.

Next, we inferred 3D joint positions using a trained convolutional pose machine (CPM; Bala et al 2020). We used a loss function that incorporated physical constraints (such as preserving limb length, and temporal smoothness) to refine joint localization. We found residual variability in limb length across subjects after reconstruction, between subjects, particularly for the arm, resulting in poses that were highly specific to individual subjects. To prevent subject-specific limb lengths from biasing subsequent behavior identification, we augmented the original 13 inferred landmarks to include two new ones (positions of left and right elbows) using a supplementary trained CPM model (method described in Bala et al., 2021). Thus, the augmented reconstruction resulted in 15 annotated landmarks for each image.

### Pose preprocessing

To discover poses, we applied a number of smoothing and transformation steps to the 3D pose data. First, we transformed the reconstructed space to a reference space that was measured using the Optitrack system (Bala et al 2020). Then, we ignored any frame where a limb was outside the bounds of the cage due to poor reconstruction, or residual frames where subject poses were still subject to collapse (defined as where the mean limb length < 10 cm). Next, we interpolated over any segments of missing data (lasting at most 10 frames, or 0.33 sec) using a piecewise cubic interpolation. Note that only a small number of frames were removed after this procedure; specifically, 0.64% of frames on average were ignored.

We next normalized the orientation of poses on individual frames. To this end, we translated the 3D joints to a common reference point by subtracting the position of the *neck* landmark. Next, we scaled poses to all have the same size, so that the spine was of length 1.0 (arbitrary units). Finally, we rotated poses to face a common direction. To do this rotation, we first defined two vectors, one corresponding to the spine (neck to hip landmarks), and the other to the expanse of the shoulders (left and right shoulder landmarks, which was then centered on the neck landmark). Poses were then rotated such that the plane defined by these vectors faced the same direction (in essence, so that the torso faced the same direction).

We next aligned individual datasets, inspired by the *mutual nearest neighbors* procedure developed for correcting for batch effects in genomics data (Halgverdi et al., 2018). Broadly speaking, this algorithm first seeks similar poses between datasets, and then applies a locally linear correction to align similar poses. Specifically, for two datasets **X1** and **X2**, we first performed two K-nearest neighbor (KNN) searches (for samples x_2_ in **X1**, and x_1_ in **X2**) using a euclidean distance and searching for K=100 samples. On the basis of this search, for each sample, we defined a *mutual nearest neighbors* set, namely, samples from each dataset that were within each other’s nearest neighbor set. We then computed a correction vector *c* for each sample in **X2** as the mean of the difference between the sample and its mutual nearest neighbors, weighted by their distance. Samples that had no mutual nearest neighbors did not have a correction vector computed. As poses vary continuously in time, we then used a median filter (15th order) to smooth out the correction vectors in time, obtaining a correction matric **C**.The aligned dataset **X2’** was defined:

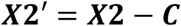

### Feature engineering

To label pose samples, we first defined a set of 23 features derived from the preprocessed pose data. The first 19 features were the angles at each joint (i.e. the vertex of each triplet of adjacent landmarks). The other four features were (1) the overall speed of the subject (calculated from the centre-of-mass), and (2,3, and 4) the speed of the subject in the three canonical dimensions (X, Y, and Z). To prevent feature bias due to differences in scale during embedding, we normalized each set of features (joint angles, COM velocity, and planar velocities) to the range [0 1]. This was performed independently for each subject.

We then concatenated data from all 18 datasets. To mitigate possible effects of noise, we applied a Principal Component Analysis (PCA), and extracted the first 16 PCs (which accounted for 95% of the explained variance). Projections onto these PCs served as the features that were then used in subsequent embedding and clustering.

### Posture identification via embedding and clustering

We created behavioral maps by embedding the extracted pose features into two dimensions using Uniform Manifold Approximation and Projection (UMAP; McInnes et al., 2014), using a euclidean distance metric. We set the parameters *min_dist*=0.001 and *n_neighbors*=20, which we found to be a good balance between separating dissimilar poses, while combining similar ones.

To define behavioral clusters, we first estimated the probability density at 200 equally interspersed points both in the first and second UMAP dimensions. This produced a smoothed map of the pose embeddings, with clearly visible peaks. We then employed the *watershed algorithm* on the inverse of this smoothed map (Berman et al., 2014). This algorithm defines borders between separate valleys in the (inverse of) the embedding space. Thus, the algorithm determines sections of the embedding space with clearly delineated boundaries (i.e. clusters). Samples were then assigned a *posture* label according to where they fell within these borders.

### Transition probabilities

We defined transition matrices between postures. Specifically, the transition matrix *M* for a transition lag of *T* was defined as:

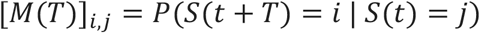

Which describes the probability that the subject would go to posture *S=i* given posture *S=j* at time *t* after *T* transitions. Note that we did this for transitions between different postures (thus, for a transition lag T=1, it is impossible for a posture to transition to itself). We performed this for each dataset individually. The resulting transition probability matrix is a directed graph, where nodes are the individual postures, and the probabilities are the weights on the edges between nodes. This formalization allows us to leverage tools from graph theoretic work.

### Measures of behavioral organization

To discover how postures are organized, we employed a hierarchical clustering algorithm named Paris (Bonald et al., 2018), using the *sknetwork* library (https://scikit-network.readthedocs.io/). This algorithm employs a distance metric based on the probability of sampling node pairs and performs agglomerative clustering. Paris requires no user-defined parameters (as opposed to another popular graph clustering algorithm, Louvain, which can perform hierarchical clustering according to a user-supplied resolution parameter). It is equivalent to a multi-resolution version of the Louvain algorithm (Bonald et al., 2018). The result of this algorithm is a dendrogram describing the relation between different posture transitions (which we will refer to as the behavioral dendrogram). To segment pose transitions into modules, we determined the *modularity score* (see below) for different cuts of each dendrogram (space equally from 0.4 to 1.4). We then determined module assignment by cutting the behavioral dendrogram where the modularity score was maximized.

We leveraged three important graph-theoretic metrics to assess behavioral composition:

- *Modularity Score*: The modularity score describes the degree to which postures transition within, rather than between, modules. Transition probability matrices with high modularity scores exhibit a high probability of transitions within modules, but not between modules. Modularity was calculated with the matlab function “modularity.m”.
- *Dasgputa Score*: To assess whether the graph defined by posture transitions truly reflected hierarchical organization, we calculated the *Dasgputa Score (*Dasgupta, 2016). The Dasgupta Score is a normalized version of the Dasgupta Cost, which defines the cost of constructing a particular dendrogram, given distances between nodes. The Dasgupta Score thus provides quantification of the quality of the hierarchical clustering. We calculated this score using the function “dasgupta_score” in the *sknetwork* library.
- *Adjusted Mutual Information Score*: We assessed whether modules were composed of similar poses using the *Adjusted Mutual Information Score (AMI;* Vinh et al 2010). This measure assesses the information (in bits) about one set of cluster assignments given knowledge of another. It is *adjusted* because given two random clusterings, the Mutual Information score is biased by the number of clusters; the adjustment thus corrects for this bias. AMI was computed using the Matlab function “ami.m”.

We further leveraged these measures to determine the timescale of behavioral organization, and make subsequent comparisons between datasets. Specifically, we extracted the half-life associated with the various measures as a function of transition lag. As a concrete example, we extracted the modularity score using transition matrices at different lags. We then fit an exponential model of the form:

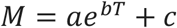

Where *M* is the modularity score at transition lag *T*. From this, we can determine the half-life of the curve *H* as:

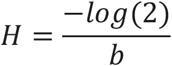

We repeated this analysis using AMI scores between consecutive lags.

### Statistical testing

For the present study, we sought to delineate how the structure of behavior changes with externally imposed task demands. Thus, we grouped sessions into *task ON (controlled depletion task or random depletion task)* or a *task OFF* (*no task sessions)*.

#### Hierarchical and Modular organization

To determine statistical significance of modular and hierarchical organization of behavior for any one dataset, we compared modularity and hierarchy to a transition matrix defined by random transitions. To this end, for each dataset, we shuffled the pose labels across the whole session, (2) re-built the transition matrix, (3) applied hierarchical clustering, and then (4) recomputed the modularity and Dasgupta scores. This was performed 100 times. The p-value was computed by comparing the random distribution to that of the observed, for each dataset individually. We repeated this analysis for transition matrices defined by different lags.

#### Module stability across transition lags

To determine if behavioral modules were similar across behavioral trees constructed from different transition lags, we compared module clusterings of consecutive lags (i.e. at lag t+T and t+T+1). To assess statistical significance, we again used a randomization approach. We randomized module labels, and then recomputed the AMI. This was performed 100 times, from which we determined the p-value.

#### Comparison of Hierarchical and Modular organization as a function of task and individual

To compare modularity as a function of task and individual, we performed a 2-way ANOVA on modularity scores obtained from transition matrices of lag=1. We performed the same analysis for Dasgupta scores, in order to compare hierarchical organization.

#### Module stability between datasets as a function of task and individual

We compared the composition of postures into modules across datasets, as a function of task and individual. To this end, we computed the AMI between module assignments (determined from a transition matrix of lag T=1) of all (unique) pairs of datasets.

To determine if a particular combination of task/individual exhibited stable module assignments, we employed a randomization procedure. Namely, we randomized the module labels for any one dataset, recomputed the AMI, and repeated this 100 times in order to get a random distribution. P-values were determined by comparing the observed AMI to the random distribution.

To determine if task/individual affected stability between paired datasets, we used a 2-way ANOVA. The first factor had 3 levels corresponding to individual pairings (subject C-Y, C-C, and Y-Y). The second factor also had 3 levels (task OFF-OFF, ON-ON, and OFF-ON).

To determine if within-subject modules were more stable than between-subject modules, we first collapsed dataset pairs into within-subject (Subjects C-C and Y-Y) and across-subject (Subjects C-Y) groups. Then, because we found AMI varied by task and individual, we partialed out the effect of task by subtracting the mean AMI associated with each task pair. Significance was assessed with an unpaired T-test.

We used a similar procedure to compare the effect of environmental demands. Namely, we considered two groups of dataset pairs, either task OFF-OFF or task ON-ON, and partialed out the effect of individuals by subtracting the mean AMI associated with individuals. Significance was assessed with an unpaired T-test.

#### Comparison of behavioral organization timescales as a function of task and individual

To compare the timescale across which behavior is organized, we compared half-lives of modularity, and AMI curves (i.e. either of these measures as a function of transition lag). We then performed a 2-way ANOVA on half-lives, with individual and task as the factors.

## Acknowledgements

We thank the Hayden/Zimmermann lab for valuable discussions. This research was supported by a National Institute on Mental Health grant R01 to BYH (125377), an NSF award to HSP, BYH, and JZ, and a UMN AIRP award (to BYH and JZ).

